# DNA metabarcoding revealed how time and space do matter -sex does not- in the dietary variation of the endangered Dupont’s Lark

**DOI:** 10.1101/2024.03.17.585437

**Authors:** Julia Zurdo, Daniel Bustillo-de la Rosa, Adrián Barrero, Julia Gómez-Catasús, Margarita Reverter, Cristian Pérez-Granados, Jesús T. García, Javier Viñuela, Julio C. Domínguez, Manuel B. Morales, Juan Traba

## Abstract

A species’ diet is highly dependent on the availability of food resources in space and time, as well as on intrinsic factors such as sex or age. Accurate assessments of variations in the diet composition of bird populations across spatial scales, seasons and demographic groups are essential not only for understanding the basic ecology of species, but also for the conservation of endangered ones. However, our current knowledge about how birds’ diet change according to spatio-temporal variations or intrinsic factors is very limited. Here, we used a multi-marker metabarcoding approach to characterize the diet of a declining shrub-steppe passerine, the Dupont’s Lark (*Chersophilus duponti*), throughout a large part of its global distribution range. We also investigated spatial, phenological and sexual variations in its diet. Using markers from two genomic regions (18S and COI), we analyzed fecal samples from 303 adult Dupont’s larks from Morocco and Spain during the breeding and non-breeding seasons. Overall, arthropods from the orders Coleoptera, Lepidoptera, Julida and Orthoptera were the main prey consumed by Dupont’s Larks. We found that Dupont’s Lark diet varied spatially, as well as temporally, reflecting dietary plasticity in response to changes in prey availability across landscapes and the species’ phenological periods. High dietary overlap and no differences between sexes were observed, suggesting similar foraging behavior and nutritional requirements in both sexes. This is the first study providing detailed information on Dupont’s Lark food ecology over much of its distribution, which is fundamental for the management and conservation of this declining steppe species.

## Introduction

The study of animals’ diet is central to understand the survival and maintenance of individuals and populations [1], as diet influences many dimensions of animals’ life history, including physiology, habitat use, migration, survival and breeding success [2]. However, accurately characterizing animal diets is challenging, because of the effort required to collect precise data and the variations in food resources used by animals, which can be affected by both abiotic and biotic factors [3, 4].

The diet of a species can change over time and space owing to variations in availability of food resources [5]. Arthropods, for instance, vary in abundance throughout the year, and they typically have an uneven spatial distribution, even at local scale (i.e., within habitats [6]). Consequently, seasonal, annual, and regional variations may occur in the diet of insectivorous animals [2, 7, 8]. Accordingly, several insectivorous animals exhibit dietary plasticity, which is a key factor that enables species to deal with environmental changes affecting prey distribution and abundance [9]. According to optimal foraging theory, the diet of generalist predators is expected to vary seasonally, shifting between alternative prey species depending on which are more abundant [10]. It is therefore essential to investigate dietary variation, at both spatial and temporal scales, to fully understand the food ecology of animals, which is, in turn, critical for wildlife management and the conservation of endangered species [11]. Such knowledge can help identify when these species are more vulnerable or when their populations are most likely to experience food limitations [12].

Diet can also vary among individuals of the same population due to intrinsic factors, such as sex, reproductive status or age [8, 13, 14]. Indeed, sex-related differences on birds’ diet have been typically attributed to morphological and behavioral differences [15, 16], but also to differing nutritional demands during the breeding season due to distinct reproductive roles [17]. Sexual dimorphism is expected to decrease dietary overlap and intraspecific competition for food resources [18, 19], although dietary differences between sexes have been also found in species with reduced dimorphism (e.g., [16]). Altogether, an accurate knowledge of dietary variation at both species-specific and spatio-temporal levels might be crucial in identifying the underlying causes of decline of endangered species.

Dupont’s Lark (*Chersophilus duponti*) (Fig 1) is a scarce steppe passerine of the Alaudidae family, whose distribution is restricted to Spain in Europe, and to the Maghreb (Morocco, Algeria and Tunisia), Libya and Egypt in Africa [20]. Dupont’s Lark is listed as “Vulnerable” at a European level [21], and in Spain the species has recently been listed as “Endangered” [22]. This species shows sexual dimorphism, evidenced by the larger size of the males [23, 24], and is strongly habitat-selective, occurring exclusively in flat (slope less than 15%), treeless shrub-steppe systems [25], which makes it highly vulnerable to habitat alterations. In fact, the land-use changes and habitat fragmentation that drive the regression and degradation of steppes in Spain and the Maghreb [20, 26–28] affect the entire passerine community which depend on this particular habitat [29], and are leading to the loss and isolation of Dupont’s Lark populations [30]. Indeed, the European Dupont’s Lark population showed a decrease of 29.9% from 2004 to 2022 [31]. In Morocco, the species has also experienced a population decline and a reduction of its breeding area [20]. Population sizes and trends in the rest of its African distribution are less well known, but a recent study suggests regressive trends in the Maghreb, at least in Tunisia [28].

**Fig 1.**
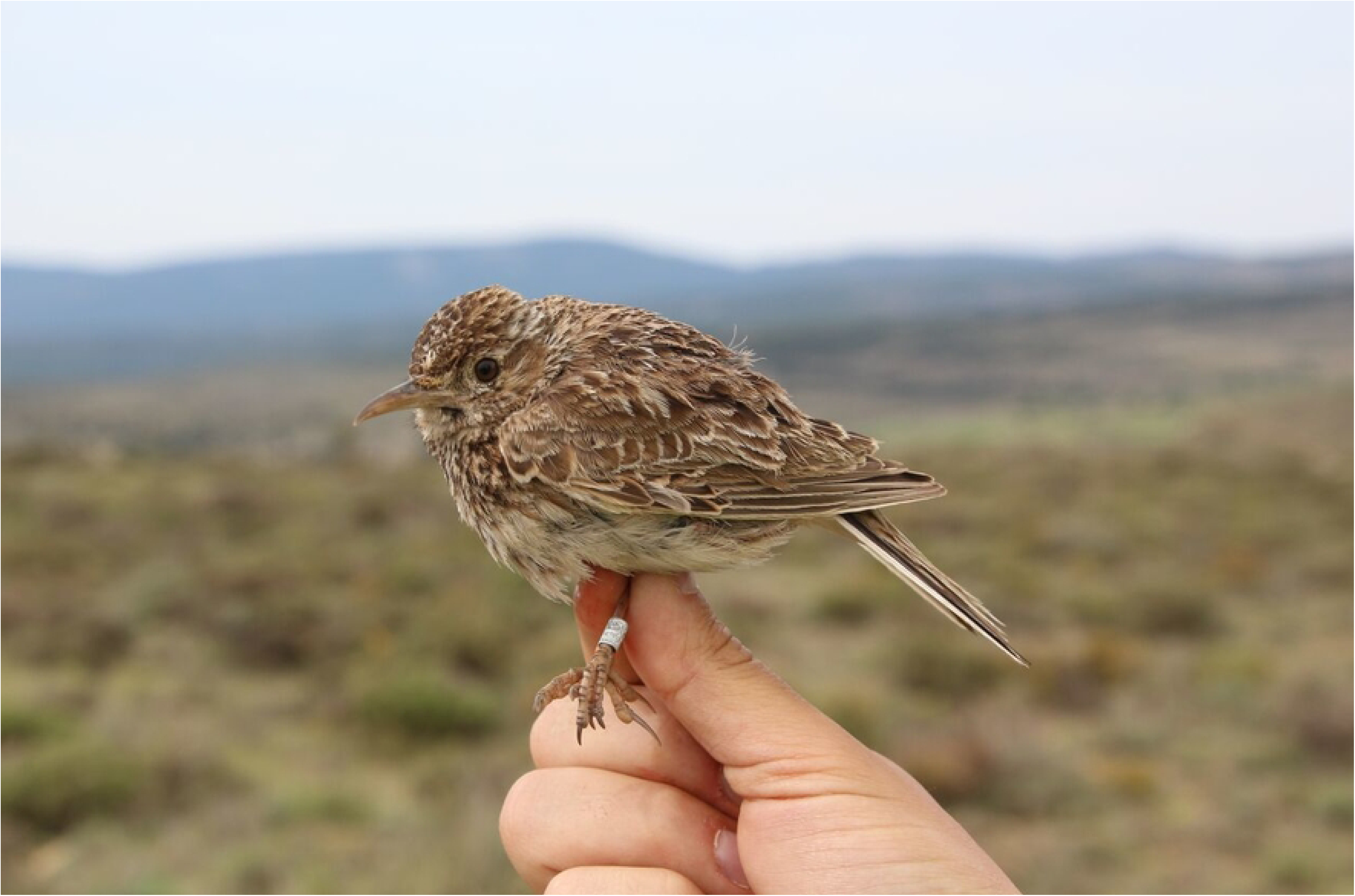
Cover photograph. A male Dupont’s Lark (*Chersophilus duponti*). Photographed by Adrián Barrero – TEG-UAM.

Knowledge on Dupont’s Lark diet is limited and restricted to specific Spanish populations. [32] described the species as insectivore and occasionally granivore, which was supported by a study based on visual examination of four gizzard contents collected in early autumn [33]. The diet of nestlings of this species was studied by [34], who noted the importance of Orthoptera, Lepidoptera, Araneae and Coleoptera. Recent studies using DNA metabarcoding have provided the first detailed information on adult and nestling diet of this species [14, 35, 36], but the information in these studies is limited to data collected from a single study area in Central Spain during the breeding season. These local studies revealed that Dupont’s Lark most frequently consumed Coleoptera, Lycosidae (Araneae), Julidae (Julida), Orthoptera, Cydnidae (Hemiptera) and Lepidoptera [14], showing age-related dietary differences [35]. However, there are currently no studies assessing how the diet of this specialist species varies over large spatial scales and across different periods of the species’ phenology.

In this study, we used multi-marker DNA metabarcoding to conduct the first comprehensive diet analysis of Dupont’s Lark over most of the range of the nominal *C. d. duponti* subspecies [20]. DNA metabarcoding is a molecular technique that overcomes the limitations of traditional methods (e.g., time-consuming, coarse taxonomic resolution, lack of identification of prey items that leave no hard remains [37, 38], providing the ability to identify a wider range of ingested taxa with higher taxonomic resolution [39]. Here, we aimed to provide an accurate description of dietary composition across spatial scales, phenology and demographic groups, and to explore whether the diet of Dupont’s Lark varies and what factors determine such variation. We predicted that the composition of Dupont’s Lark diet will vary at both macro-regional (countries) and regional scales (within Spain), as well as at different periods of the species’ phenology, likely reflecting spatial and temporal changes in prey availability. We also expected sexual differences in diet composition, probably derived from the morphological differences between sexes [24] and reflecting different nutritional demands of both groups during the breeding season. The findings from this study might be useful to inform Dupont’s Lark conservation strategies across its distribution range to ensure the survival of this species, which could also benefit other declining steppe passerines.

## Materials and methods

### Study sites

Fieldwork was carried out in two countries, Spain and Morocco, covering a large part of the species’ distribution range [40] (Fig 2).

**Fig 2.**
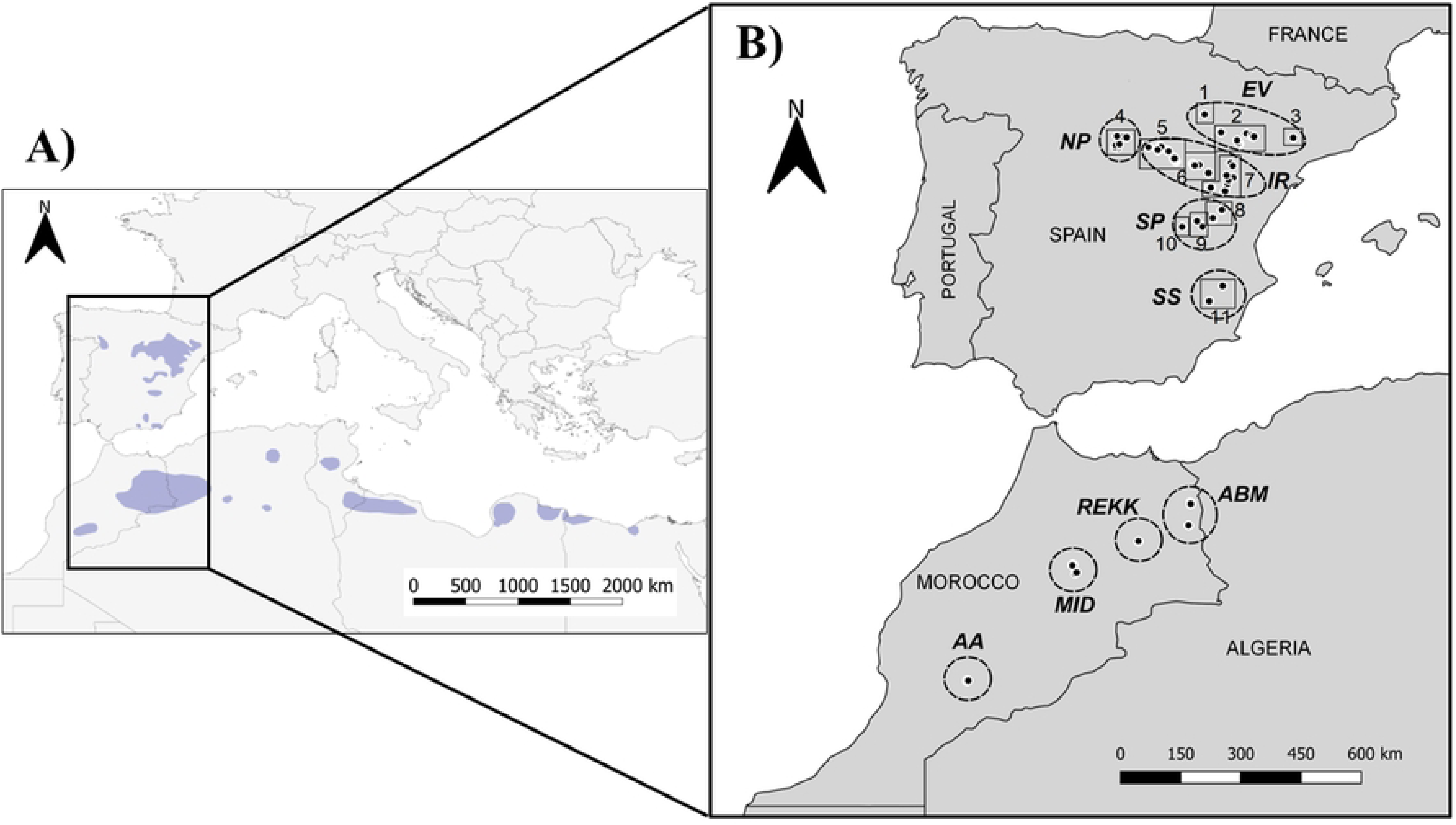
Location of the study area. (A) Location of the study area within the Dupont’s Lark global distribution (blue polygons) based on [21]. The subspecies *Chersophilus duponti duponti* is distributed in the Iberian Peninsula, Morocco, northern Algeria and Tunisia; the subspecies *C. duponti margaritae* is distributed in southern Algeria and Tunisia, Libya and Egypt. (B) Spanish and Moroccan sampling points (black dots). Fieldwork regions are indicated with dashed lines: Northern Plateau (NP), Ebro Valley (EV), Iberian Range (IR), Southern Plateau (SP), and Southern Spain (SS) in Spain; Aïn Bni Mathar (ABM), Plateau of Rekkam (REKK), Midelt-Missour (MID), and Anti-Atlas (AA) in Morocco. Solid lines and numbers from 1 to 11 denote Spanish study sites defined in the study: 1. Ablitas; 2. Ebro Valley; 3. Alfés; 4. Hoces y Corcos; 5. Western Iberian Range; 6. Eastern Iberian Range; 7. Altiplano; 8. Ademuz; 9. Carboneras de Guadazaón – Cardenete; 10. Valeria; 11. Cieza.

In Spain, sampling was conducted in spring, during the Dupont’s Lark breeding seasons (March-June) of 2017-2019 and in autumn, during the non-breeding season (September-November) of 2017. We collected fecal samples in 11 study sites distributed over five broad geographic regions throughout the range (see regions description in [41]) (Table 1, Fig 2): Northern Plateau (NP), Ebro Valley (EV), Iberian Range (IR), Southern Plateau (SP), and Southern Spain (SS). IR and EV regions are the main core areas of Dupont’s Lark, where the largest patches of continuous typical habitat of the species in Spain are found, whereas the rest are isolated and highly fragmented areas [42]. The dominant landscape, in all regions, is flat steppes covered by low scrub and a high proportion of bare ground [25, 40].

**Table 1.**
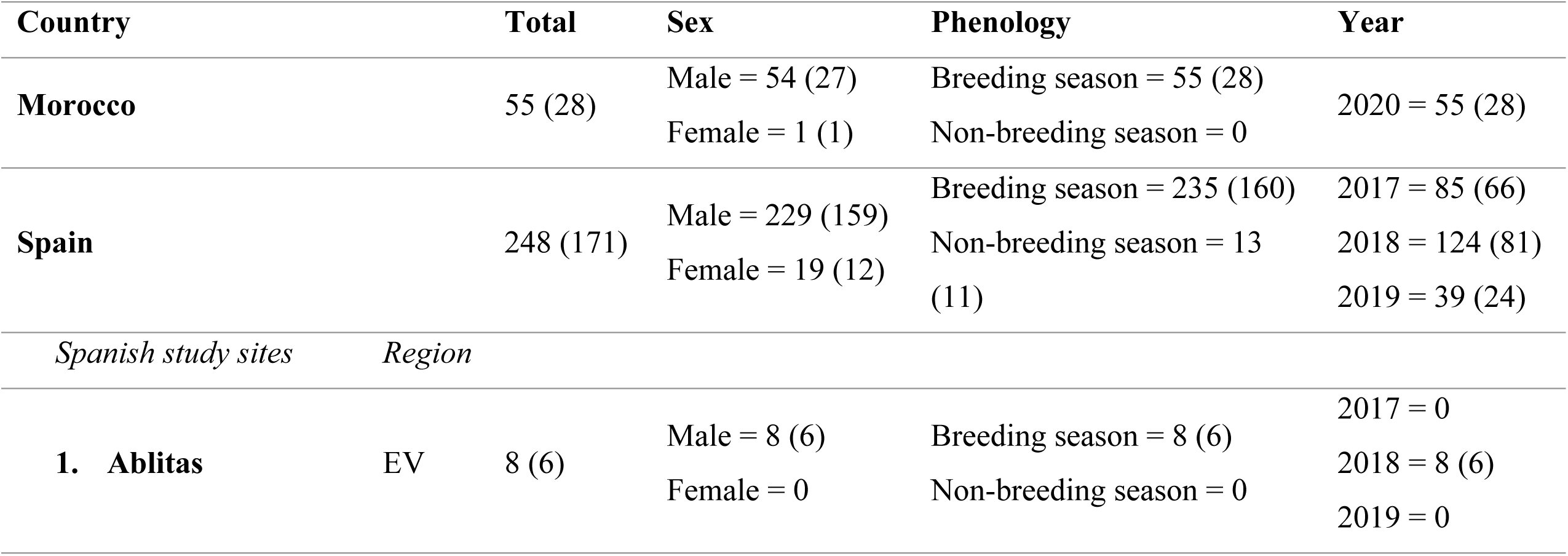
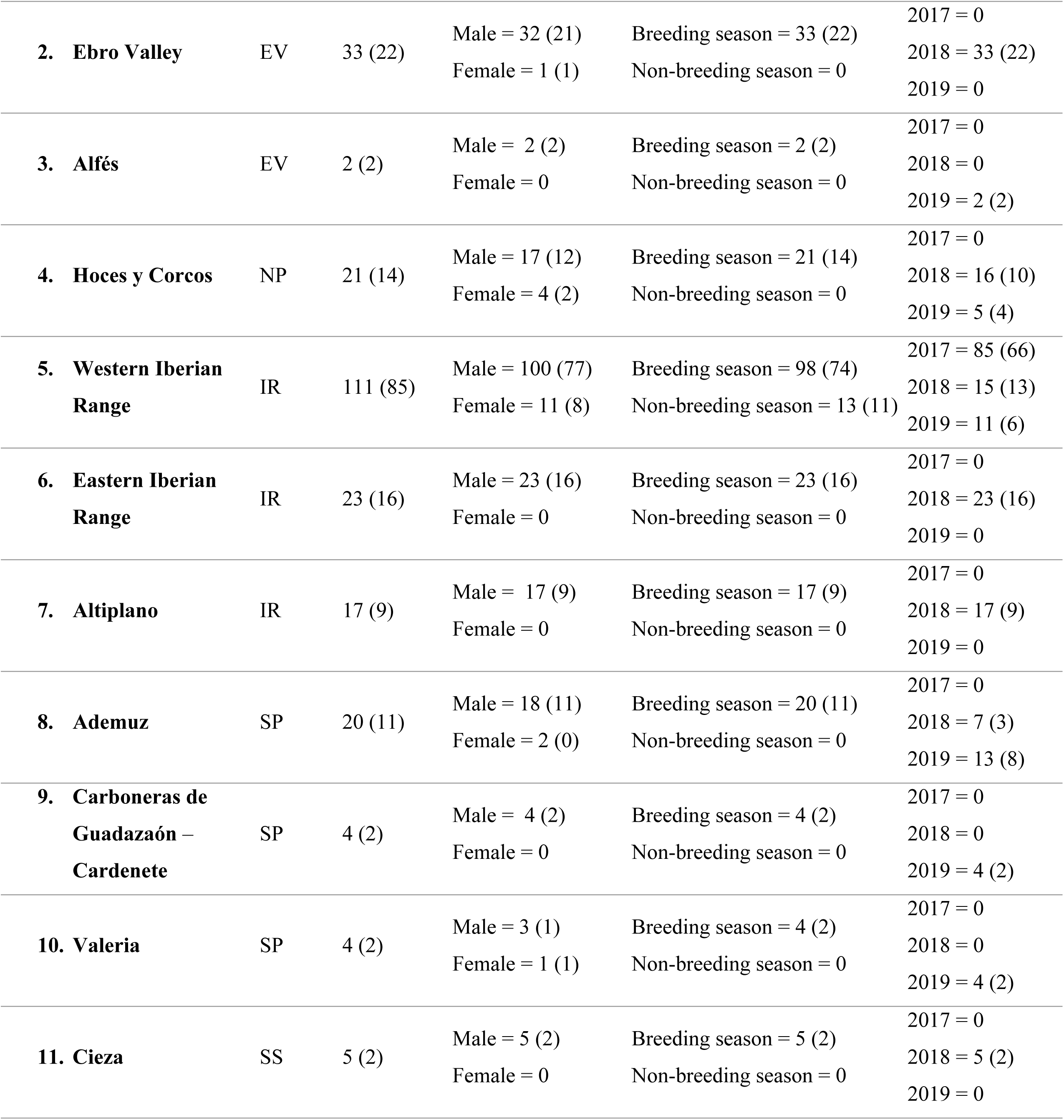
Number of Dupont’s Lark samples collected in Morocco and Spain and across study sites of Spain (numbers from 1 to 11 in **Fig 2B**), broken down by sex, species’ phenology, and year. Successful fecal samples after molecular analysis are indicated in brackets. The region to which each Spanish population belongs is specified: Ebro Valley (EV), Northern Plateau (NP), Iberian Range (IR), Southern Plateau (SP), and Southern Spain (SS).

In Morocco, we sampled in late winter/early spring (February-March) of 2020, coinciding with the breeding season of larks in such area. We collected samples throughout the range, in the four geographic regions determined by [20] as suitable sites for the species: Aïn Bni Mathar (ABM) in northeastern Morocco, Plateau of Rekkam (REKK) and Midelt-Missour region (MID) at the central plateau, and the Anti-Atlas region (AA) in southwestern Morocco (Fig 2). Steppe vegetation was dominated by alfa grass *Macrochloa tennacissima* except in the southwestern (AA), where *Artemisia* spp. was the dominant species [20].

### Sample collection

For collecting Dupont’s Lark fecal samples (*n* = 303, Table 1), we first captured adult individuals using spring-traps baited with mealworms (*Tenebrio molitor*) and a species-specific recording to attract them, which is a male-biased sampling [43]. Individuals were ringed to avoid pseudoreplication and released at the site of capture. Sex was determined through wing length following [23], who determined that adult males had wing lengths > 97 mm while females had wing lengths < 97 mm. Active brood patch in females during the breeding season was also a sex discriminant trait. We collected fresh fecal samples from the ground at the spring-trap during the capture or produced in the ringing bags. To prevent contamination, in Spain the ringing bags were washed with bleach before each use, whereas in Morocco sterile filter paper placed inside the bags was used. Samples were stored in individual 1.5-mL plastic vial tubes with 98% ethanol and refrigerated at -20 °C until processed in the laboratory. All birds were captured and handled in accordance with both national and international guidelines and under permits from Moroccan and Spanish authorities. All procedures were approved by the Local Ethical Committee for Animal Experiments of the Universidad Autónoma de Madrid (CEI80-1468-A229).

### Molecular sample processing and bioinformatics

The QIAamp PowerFecal DNA Kit (Qiagen) was used to extract DNA from fecal material, following the manufacturer’s instructions. Before extraction, ethanol was removed from the samples by decanting following 30 min of centrifugation at 12,000 rpm and heated at 50 °C until the ethanol was vaporized. DNA extraction was performed by the Genomics and NGS Core Facility at the Centro de Biología Molecular Severo Ochoa (CBMSO, CSIC-UAM, Spain). To amplify prey DNA, we used two marker sets: a universal eukaryote 18S marker (miniB18S_81 [44]), and the arthropod marker ZBJ [45] for COI region. Each marker was amplified in an independent PCR reaction, in which a negative PCR control (DNA-free) was included. Products were subjected to a second PCR to perform indexing and attach Illumina sequencing adaptors. PCR reactions were conducted following [44] and [45] for each marker, respectively. PCR products were purified using AMPure XP beads (Beckman Coulter) and checked in Bioanalyzer before pooling per marker in equimolar amounts. The final two libraries were sequenced in an Illumina MiSeq NGS platform using a v3 MiSeq Reagent kit, following the manufacturer’s instructions. Amplification, library preparation and sequencing were carried out by the Genomics Unit of the Fundación Parque Científico de Madrid (Spain).

Bioinformatic processing of sequencing reads was performed using MJOLNIR pipeline (Metabarcoding Joining Obitools and Linkage Networks In R; pipeline steps in Appendix S1), with a separate analysis for each molecular marker. We first used OBITools [46] to quality filter and align paired-end Illumina sequences, and then we implemented VSEARCH [47] to remove chimaeras. We clustered the sequences into molecular operational taxonomic units (MOTUs) using swarm [48], which is based on an iterative aggregation of sequences that differ less than a given distance. For the taxonomic assignment, we first created a reference database in ecoPCR format for each molecular marker, obtained from the download of all 18S and ZBJ sequences from NCBI database.

We then used ecotag from OBITools to match the MOTUs generated to the reference sequences. We finally removed pseudogenes using LULU [49]. MOTUs were identified with the most resolved taxonomic assignment possible, and those identified to species or genus were manually confirmed using the BLAST algorithm (NCBI). We removed every taxa not belonging to Animal kingdom, as well as mammals (human), birds and internal parasites (phyla Nematoda and Platyhelminthes). We also excluded taxa not considered as potential prey items (mites, ticks, springtails [14, 38]). We finally removed MOTUs representing less than 1% of the total number of dietary reads [50] to avoid incorporating false positives resulting from tag-jumping events, and samples with less than 100 dietary reads as they were considered to have failed (negative PCR controls and some Dupont’s Lark samples).

Finally, for each sample, the dietary information derived from the two molecular markers was combined using a python 3.0 script [38]. The script takes into account the differences in taxonomic resolution provided by the different markers, assuming that a dietary component obtained at a lower taxonomic resolution (e.g., order or family) by one of the marker is the same as a component of the same taxonomic group obtained at a higher resolution by the other marker (e.g., genus or species).

### Statistical analysis

For all statistical analyses, we used presence/absence data at order and family levels instead of read count, due to inherent biases present throughout the metabarcoding workflow, including differential DNA extraction success and PCR amplification rates between taxa detected within the diet [51]. All statistical analyses were carried out in R version 4.3.0 [52]. Due to the small sample size from some Spanish study sites (Table 1), those located a short distance apart (maximum distance of 85 km) and with similar ecological and geographic characteristics were merged for the analyses (hereafter referred to as ‘populations’; Table 2): study sites 2 and 3 constituted the population Ebro Valley – Alfés; and study sites 8, 9, and 10 formed the population Ademuz – Valeria. The study site number 11 (Table 1) was excluded from statistical analyses due to non-compliance with the above criteria.

**Table 2.** Populations created for statistical analyses due to the small sample size of some study sites after molecular analysis. The study sites that constitute them (numbers from 1 to 11 in Fig 2), the region to which they belong, and the number of fecal samples collected (successful samples after molecular analysis in brackets), are indicated. Region: Ebro Valley (EV), Northern Plateau (NP), Iberian Range (IR), Southern Plateau (SP), Southern Spain (SS).

| Spanish population | Spanish study site | Region | Samples |
| --- | --- | --- | --- |
| 1. Ablitas | 1. Ablitas | EV | 8 (6) |
| 2 – 3. Ebro Valley – Alfés | 2. Ebro Valley<br>3. Alfés | EV | 35 (24) |
| 4. Hoces y Corcos | 4. Hoces y Corcos | NP | 21 (14) |
| 5. Western Iberian Range | 5. Western Iberian Range | IR | 111 (85) |
| 6. Eastern Iberian Range | 6. Eastern Iberian Range | IR | 23 (16) |
| 7. Altiplano | 7. Altiplano | IR | 17 (9) |
| 8 – 10. Ademuz – Valeria | 8. Ademuz<br>9. Carboneras de Guadazaón<br>– Cardenete<br>10. Valeria | SP | 28 (15) |

To evaluate the most prevalent prey taxa within Dupont’s Lark diet, we calculated the frequency of occurrence (FOO) at family and order levels, defined as the number of fecal samples in which a family or order was detected divided by the total number of samples. We computed FOO by country, and in the case of Spain, we included population, phenology and sex groups. FOO was not calculated for Moroccan study sites due to their small sample sizes.

In order to test whether Dupont’s Lark diet varies over spatial scales (macro and country-level), the species’ phenology and sex groups, we computed multivariate generalized linear models (MGLMs) using the function *manyglm* in the *mvabund* package [53]. This function fitted the presence/absence data of prey taxa at order and family levels to binomial family generalized linear models (with a “cloglog” link function). In total, eight models were fitted, with the predictor variable being country (two levels: Spain/Morocco), Spanish population (seven levels: Table 2), phenology (two levels: breeding season/non-breeding season), and sex (two levels: male/female), and response being the prey multivariate dietary dataset at family and order level (four models for each level of taxonomic classification). To stablish appropriate comparisons, in the spatial and sex models we used only the samples collected during the breeding season, since samples from the non-breeding season are only available for males from a single Spanish population (Western Iberian Range). In the phenological model, therefore, we used only the samples collected in this population in 2017. Similarly, in order not to introduce confounding effects in the model for testing sex differences, we only used samples from Spanish populations where both male and female samples are available (see Table 1). The significance of variables was determined via likelihood ratio test using the *anova.manyglm* function with Monte Carlo resampling (999 iterations) and corrected univariate *p* values for multiple testing. When necessary, we performed pairwise comparisons using the *pairwise.comp* function of *anova.manyglm*. In addition, *p* values from univariate tests were extracted using the *p.uni =“adjusted”* argument to assess if any specific prey taxa were responsible for the spatial, temporal and sexual dietary differences. We visualized dietary differences using non-metric multidimensional scaling analysis (NMDS) based on Jaccard distance of prey families via the function *metaMDS* in the *vegan* package [54].

To quantify overlap in observed Dupont’s Lark diet between countries, Spanish populations, phenological periods and sex groups, we calculated Pianka’s index of dietary overlap [55] using the *EcoSimR* package [56]. This index quantifies the degree of similarity between two diets and ranges from 0 to 1, where 1 indicates complete overlap. In *EcoSimR*, a null model simulation, based on randomization of dietary data (here based on FOO) is generated and used in a statistical comparison to test whether the observed niche overlap differs from what would be expected by chance. We used the randomization algorithm 3 (RA3), which is a permutation test recommended for niche overlap studies that reshuffles zero and non-zero values [57], and 9999 repetitions for the simulation.

Additionally, we classified Spanish Dupont’s Lark samples into five different habitat types (based on the dominant vegetation) following [58] and expert criteria: (1) dwarf cushion-shape shrublands; (2) gypsicolous shrublands; (3) alfa grass steppes; (4) thyme-gorse shrublands; and (5) thyme-rosemary shrublands. Detailed description, sample size and location of habitat types are provided in Appendix 2 of the Supplementary Material. We then performed all the analyses described above (MGLM, NMDS and dietary overlap) in order to explore dietary differences between habitats (see Appendix 2 of the Supplementary Material for further description of statistical analysis).

## Results

We successfully amplified invertebrate prey DNA from 199 out of 303 fecal samples (69% in Spain, *n* = 171; 51% in Morocco *n* = 28; Table 1). After all filtering steps, sequences obtained from the ZBJ marker were clustered into 91 dietary MOTUs, with a mean of 9619.8 ± 5850 standard deviation (SD) diet reads and 2.6 ± 1.3 SD taxa per individual, while sequences obtained from the 18S marker were clustered into 103 dietary MOTUs, with a mean of 11,956 ± 4884 diet reads and 2.6 ± 1.5 SD taxa per individual. Combined results from both metabarcoding data sets produced 623 occurrences of prey from 185 MOTUs, with a mean of 3.1 ± 1.8 SD taxa per individual. The obtained MOTUs were identified to 62 families of 12 orders of three classes from the phylum Arthropoda (Insecta, Arachnida and Diplopoda) and one class from the phylum Mollusca (Gastropoda), with 37% of MOTUs identified to species level and 55% to genus (Table I-S3, Appendix S3). During the breeding season, the most frequent prey order in the diet of Dupont’s Lark (considering both Spanish and Moroccan samples) was Coleoptera (FOO: 60.3%), followed by Lepidoptera (40.7%), Julida (33.2%) and Orthoptera (29.2%). The families Julidae (33.2%), Noctuidae (29.2%), Acrididae (28.6%) and Tenebrionidae (26.1%) dominated Dupont’s Lark diet.

### Variation in diet composition at macro-spatial scale

MGLMs revealed that Dupont’s Lark diet during the breeding season differed significantly between countries (i.e., continents) at both order (LRT Deviance = 30.2, *p* = 0.004) and family levels (LRT Deviance = 102.3, *p* = 0.001). Univariate tests showed that differences at the order level were due to the order Orthoptera (Table I-S5, Appendix S5), which was detected with higher frequency in Moroccan samples (Fig 3). The families Acrididae and Geometridae were the main drivers of the differences at the family level (Table I-S5, Appendix S5), with the first family being consumed more frequently by Moroccan Dupont’s larks and the second family by Spanish ones (Table I-S4, Appendix S4). Despite these differences, niche overlap analysis showed a greater overlap than expected by chance (*p* < 0.001), with a value of Pianka’s index of 0.77 for arthropod families, consistent with the diet overlap observed in the NMDS (Fig 4).

**Fig 3.**
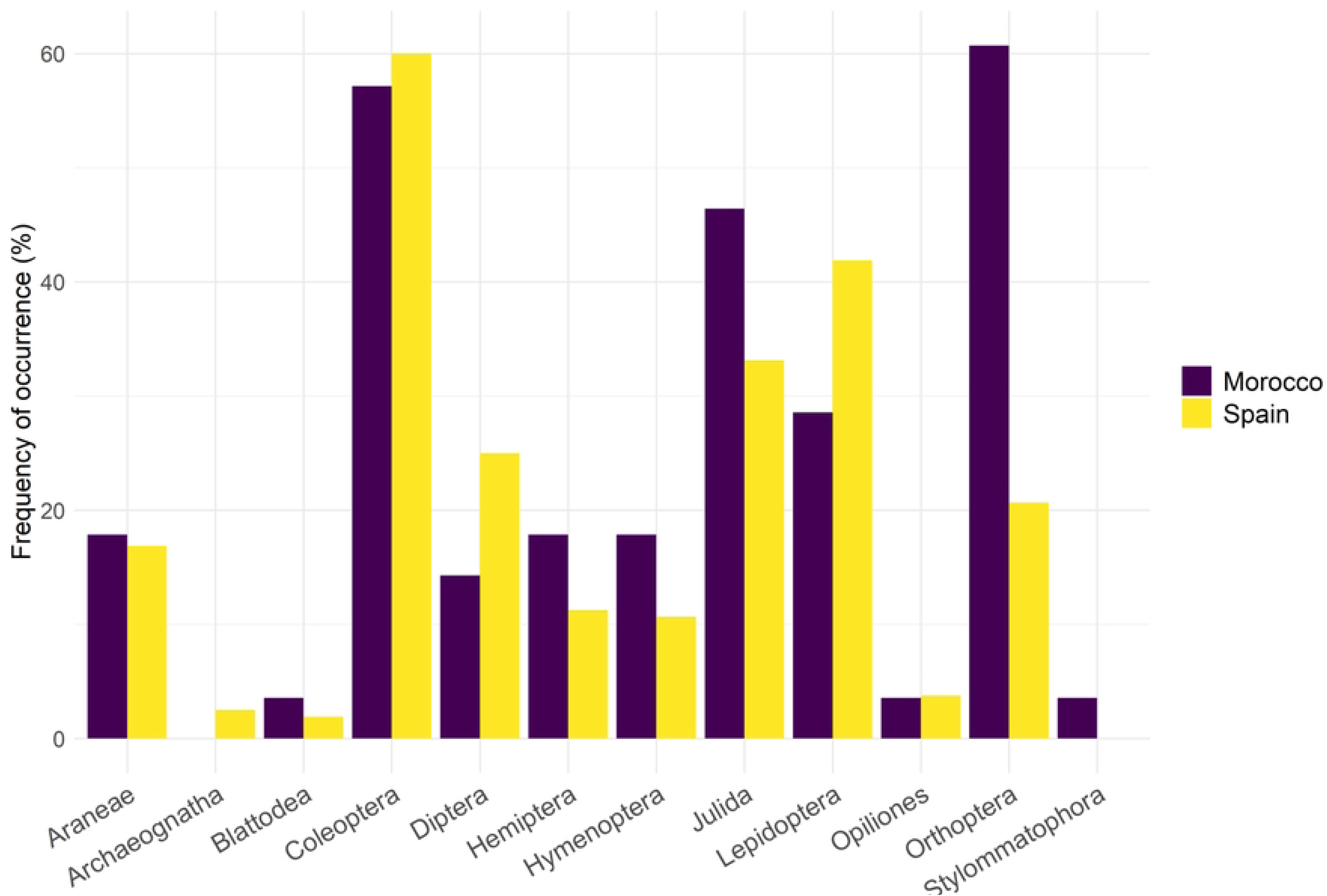
Frequency of occurrence (%) of prey orders in the diet of Dupont’s Lark during the breeding season identified using DNA metabarcoding on Moroccan (*n* = 28) and Spanish (*n* = 160) fecal samples.

**Fig 4.**
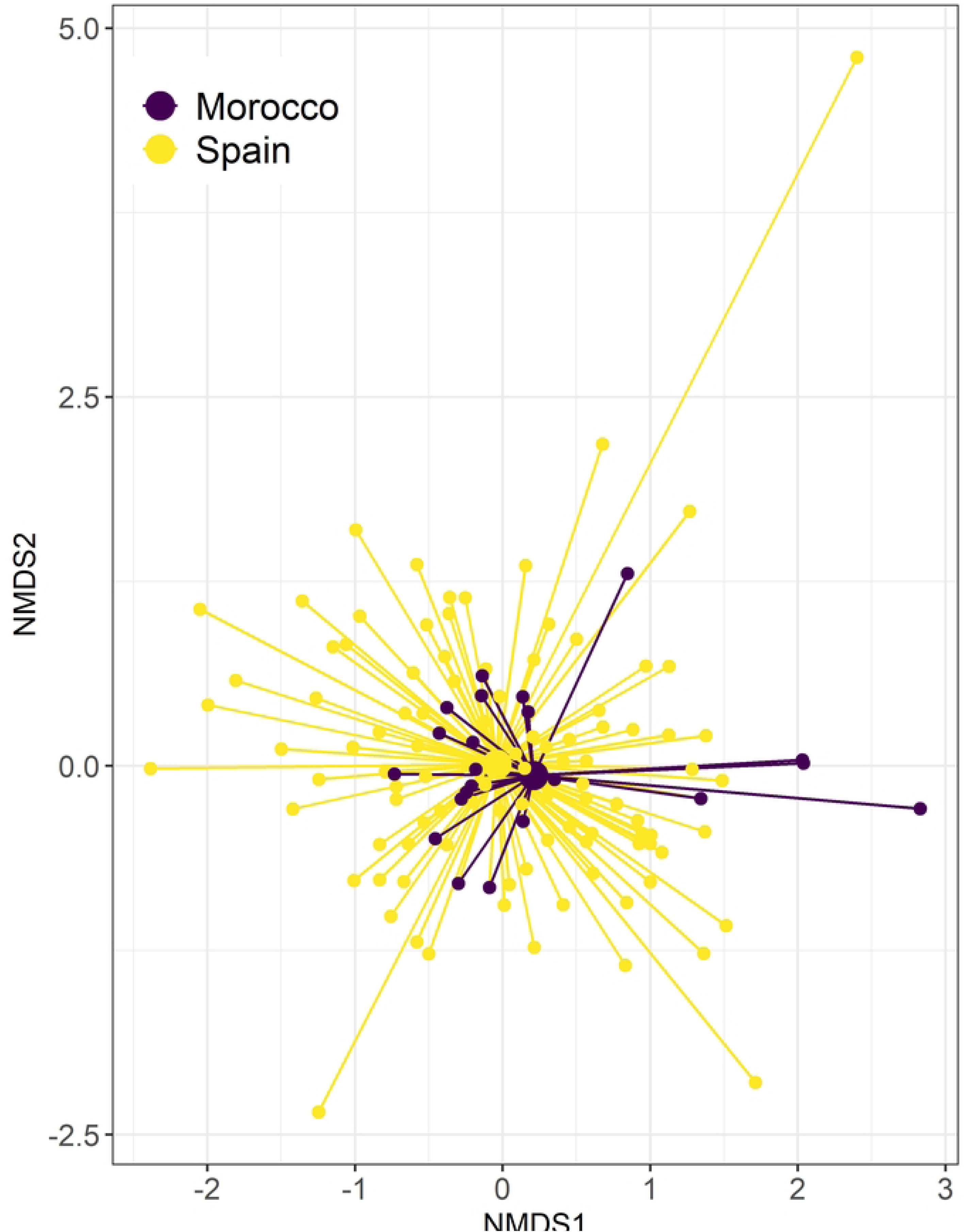
Spider plot of the non-metric multidimensional scaling (NMDS; k = 2, stress = 0.103) for arthropod prey families consumed by Dupont’s Lark in Morocco and Spain.

Smaller nodes represent individual birds with connecting lines joining the individual to the mean centroid (larger nodes) of its country.

### Variation in diet composition at country-level scale

MGLMs indicated that Dupont’s Lark diet during the breeding season differed significantly between Spanish populations at both order (LRT Deviance = 143.3, *p* = 0.001) and family levels (LRT Deviance = 345.7, *p* = 0.001). Pairwise comparisons revealed significant differences between the population 5 and the populations 2 – 3, 4 and 6 at the order level (Table II-S5, Appendix S5). These differences were due to the order Lepidoptera (Table II-S5, Appendix S5), detected with higher frequency in the samples of the population 5 than in the rest of populations (Fig 5; Table I-S4, Appendix S4). At the family level, pairwise comparisons showed significant differences between the population 5 and all other populations, with the exception of population 1, and between populations 8 – 10 and 2 – 3 (Table II-S5, Appendix S5). The lepidopteran families Noctuidae and Geometridae were significant in the univariate tests (Table II-S5, Appendix S5), with birds sampled in the population 5 showing the highest frequency for these families (59.5% and 32.4%, respectively; Table I-S4, Appendix S4). The two Dupont’s Lark individuals with dietary results from the study site 11 (not included in statistical analysis due to low sample size) consumed only prey items of the families Julidae and Tachinidae (Diptera).

**Fig 5.**
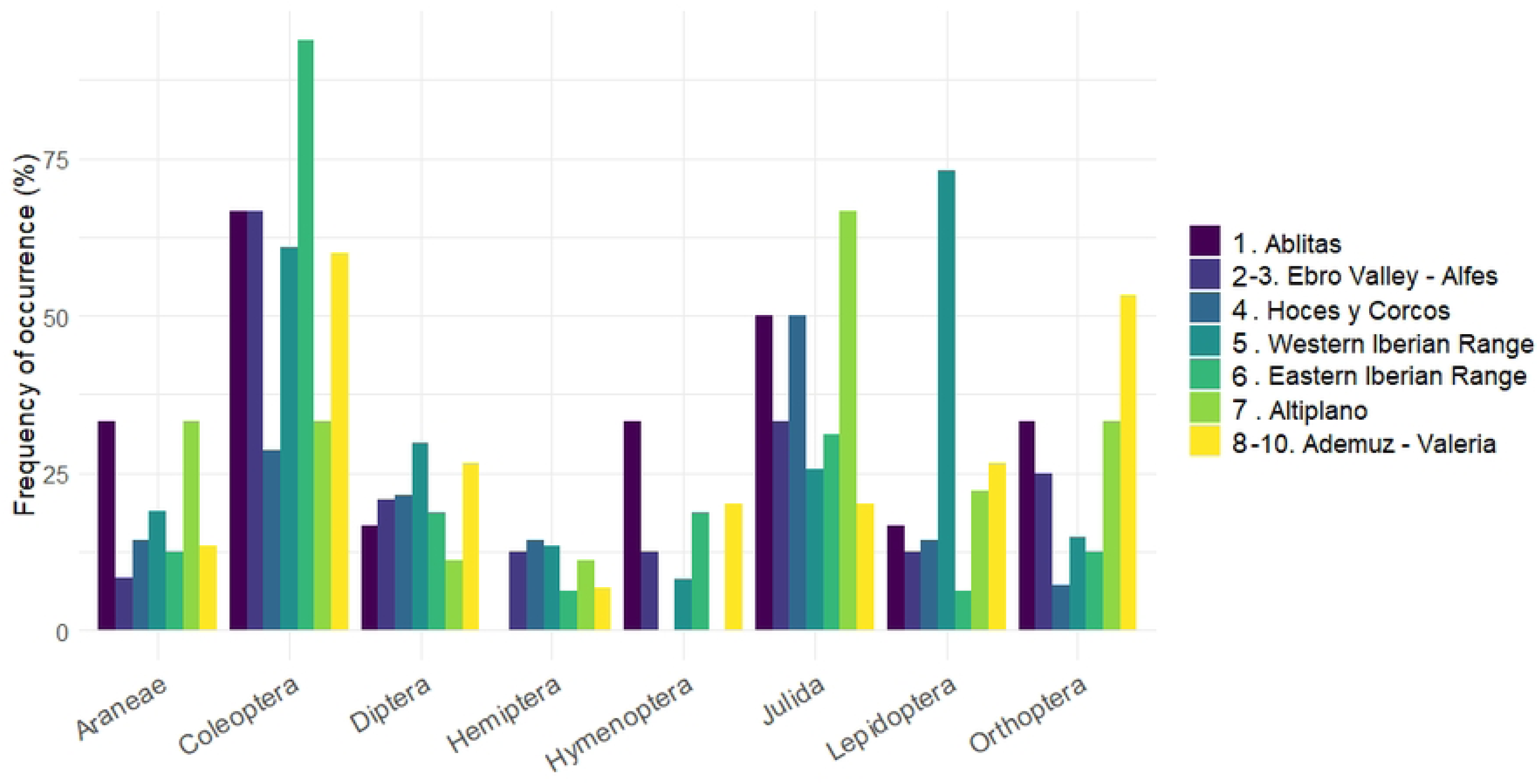
Frequency of occurrence (%) of the eight most common prey orders in the diet of Dupont’s Lark identified using DNA metabarcoding on the fecal samples from the Spanish populations sampled (numbers from 1 to 10 in **Fig 2**).

Niche overlap analysis among Spanish populations returned a mean value of Pianka’s index of 0.64, greater than the null expectation (*p* < 0.001), indicating a higher overlap in the consumption of arthropod families than expected by chance. In pairwise comparisons (Table 3) all Pianka’s index values were greater than 0.45 and statistically significant (*p* < 0.05). The lowest dietary niche overlap values were found between the population 5 and all other populations, while the highest values were observed between the population 2 – 3 and the populations 1 and 6 (0.87 and 0.86, respectively). The NMDS (Fig 6) also showed a generally high overlap between most populations, but a higher segregation was observed between the population 5 and the remaining populations.

**Table 3.**
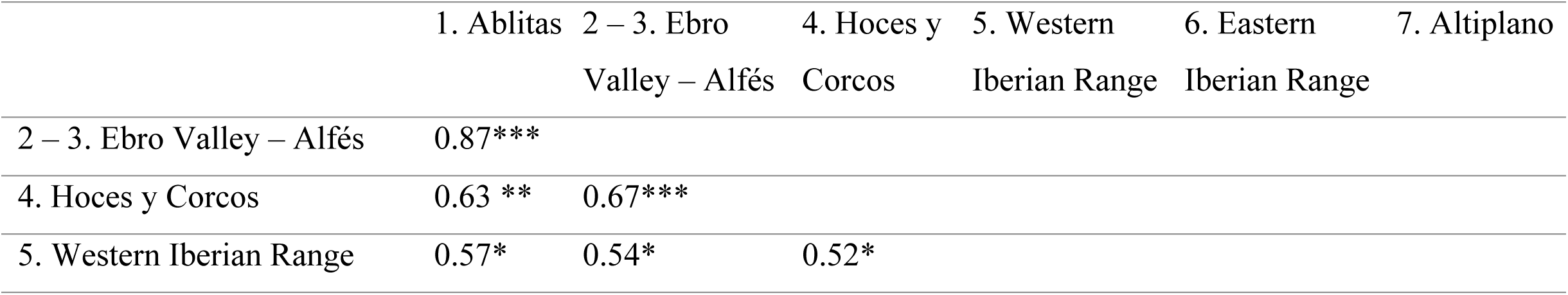
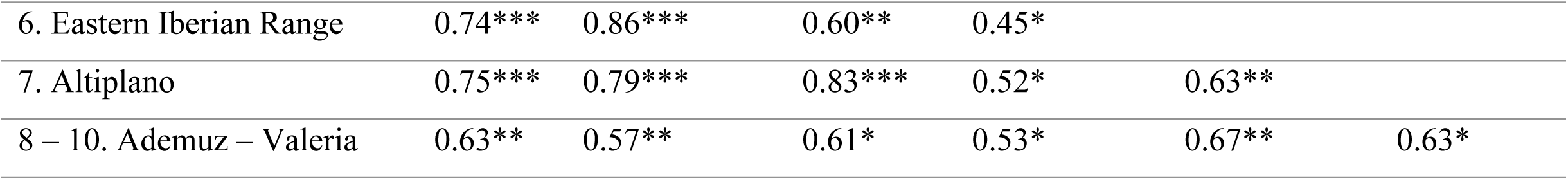
Pairwise Pianka’s index values for niche overlap estimation between Spanish populations (numbers from 1 to 10 in Fig 2) based on the frequency of arthropod families consumed by Dupont’s Lark. All values indicate statistically significant niche overlap (i.e. greater than expected by chance based on comparison with 9999 null models; * *p* = 0.05 to *p* = 0.01, ** *p* = 0.01 to *p* = 0.001, *** *p* < 0.001).

**Fig 6.**
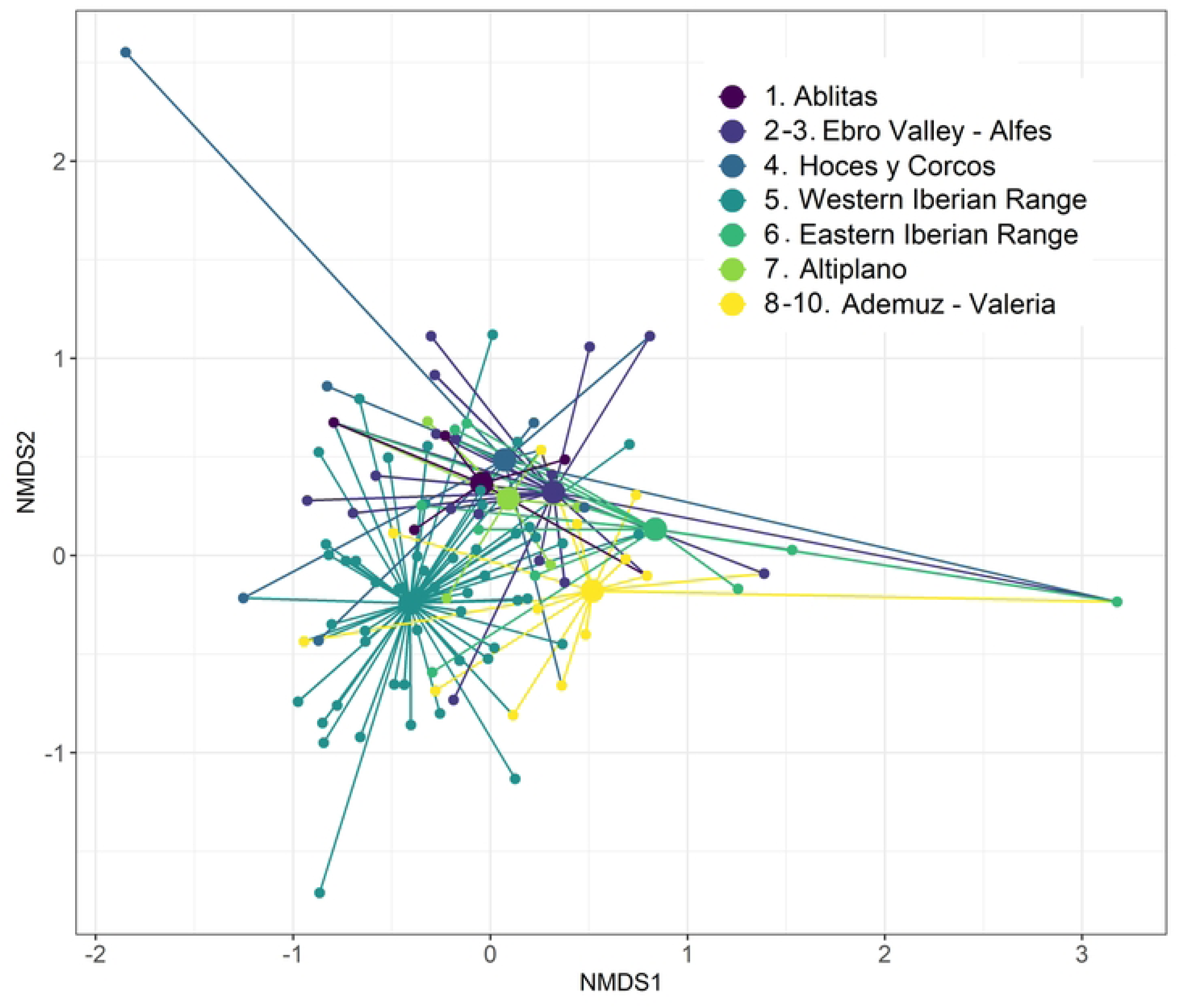
Spider plot of the non-metric multidimensional scaling (NMDS; k = 2, stress = 0.102) for arthropod prey families consumed by Dupont’s Lark across the seven Spanish populations (numbers from 1 to 10 in **Fig 2**) sampled in this study.

Smaller nodes represent individual birds with connecting lines joining the individual to the mean centroid (larger nodes) of its population.

Regarding the habitat types occupied by Dupont’s Lark in Spain, MGLMs revealed that the species’ diet during the breeding season differed significantly between them at both order (LRT Deviance = 60.9, *p* = 0.002) and family levels (LRT Deviance = 184.3, *p* = 0.001). At the order level, pairwise comparisons showed significant differences between gypsicolous shrublands and thyme-gorse shrublands (Table III-S5, Appendix S5). These differences were due to the order Lepidoptera (Table III-S5, Appendix S5), detected with the highest frequency in samples collected in thyme-gorse shrublands (Table I-S4, Appendix S4). At the family level, pairwise comparisons showed significant differences between thyme-gorse shrublands and all other habitats, and between gypsicolous shrublands and thyme-rosemary shrublands (Table III-S5, Appendix S5). The lepidopteran families Noctuidae and Geometridae, and the orthopteran family Acrididae were significant in the univariate tests (Table III-S5, Appendix S5), with birds sampled from thyme-gorse shrublands showing the highest frequency for the lepidopteran families and the lowest frequency for the family Acrididae (Table I-S4, Appendix S4).

Niche overlap analysis among habitats returned a mean value of Pianka’s index of 0.71, greater than the null expectation (*p* < 0.001), indicating a higher overlap in the consumption of arthropod families than expected by chance. In pairwise comparisons (Table II-S2, Appendix S2) all Pianka’s index values were greater than 0.58 and statistically significant (*p* < 0.05). The NMDS (Fig II-S2, Appendix S2) also showed a generally high overlap between most habitats, but a higher segregation was observed between thyme-gorse shrublands and the other habitats.

### Variation in diet composition between phenological periods

We found differences in prey composition between the breeding and non-breeding seasons within the population 5 at both order (MGLM: LRT Deviance = 30.4, *p* = 0.001) and family levels (MGLM: LRT Deviance = 54.2, *p* = 0.006). Univariate tests revealed that order Orthoptera and family Acrididae were the main drivers of the differences (Table IV-S5, Appendix S5), with both taxa being detected with higher frequency in the samples collected in the non-breeding season (72.23% vs 10.91% for both taxa; Table I-S4, Appendix S4). Despite these differences, niche overlap analysis at the family level showed greater overlap than expected by chance (*p* < 0.05), with a Pianka’s index value of 0.60. The limited diet overlap was observed in the NMDS (Fig I-S6, Appendix S6).

### Variation in diet composition between sexes

There were no differences in diet composition between sexes in the breeding season in Spain at either the order (MGLM: LRT Deviance = 6.0, p = 0.89) or family level (MGLM: LRT Deviance = 40.3, p = 0.10). A Pianka’s index value of 0.85 was obtained in the niche overlap analysis, greater than the null expectation (*p* < 0.001), and consistent with the high diet overlap observed in the NMDS (Fig II-S6, Appendix S6). Males and females consumed the orders Coleoptera, Lepidoptera and Julida with high frequency (Table I-S4, Appendix S4). Diptera was detected with higher frequency in female samples (41.7% vs 23.8%; Table I-S4, Appendix S4), as well as the Coleoptera families Carabidae and Curculionidae (Table I-S4, Appendix S4).

## Discussion

Here we used a metabarcoding approach to characterize the temporal and spatial variation of the diet of a passerine throughout most of its distribution range. More specifically, our study provides the first comprehensive comparison of dietary composition between Dupont’s Lark populations at macro-spatial (Morocco and Spain) and country-level (within Spain) scales. It also highlighted temporal differences in the diet of Dupont’s Lark across the species’ phenological periods, whereas no dietary differences related to sex were detected. Dietary variation across landscapes and periods probably reflects spatial and temporal differences in the prey community, as well as changes in prey preferences of this bird species that has been shown to exhibit diet selection [36]. This detailed dietary information is essential for understanding species’ ecology and implementing effective conservation interventions [59], especially critical for a species in such a marked population decline [31].

### Dietary composition

Dupont’s Lark can be described as a generalist insectivorous species throughout its distribution range. Larks predated on a wide range of prey groups, but most frequently fed on nutrient-rich arthropods such as beetles, lepidopterans, millipedes, and grasshoppers [60, 61]. Other important prey groups were dipterans, spiders, and hemipterans. The few previous dietary studies, based on visual examination of fecal and gizzard contents, also reported the importance of beetles and grasshoppers in the diet of Dupont’s Lark adults [33, 62], but did not observe such a high frequency of lepidopteran consumption, and did not report millipede consumption at all. In the case of Lepidoptera, this may be a consequence of the difficulties in visual identification of soft-bodied prey that leave few or no hard parts [38], which highlights the value of using molecular techniques such as metabarcoding for dietary studies of insectivorous predators. In this line, recent metabarcoding studies conducted in Central Spain on the food niche of shrub-steppe birds [14, 35] also pointed out the high frequency of consumption of beetles, grasshoppers, millipedes and spiders by Dupont’s Lark, although in the present study, which considers a larger distribution range, lepidopterans were much more often detected in the faces of the species. In addition, we detected here dietary items that had not been described in the previous studies in the diet of adults of this species, such as millipedes of the genus *Cylindroiulus*, frequently consumed in several Spanish populations, or the spider family Oxyopidae, only present in Moroccan samples. It is also remarkable the observed consumption of gastropod prey (snails) both in Spain and Morocco, but especially in the latter country, where the frequency of occurrence reached 10%. Snails represent a good source of nutrients [63] and are an abundant prey in Moroccan steppes [64]. References on snail consumption in larks are very limited (but see [65]).

### Spatial variation

Spatial dietary differences of Dupont’s Lark were observed at different scales, probably reflecting dietary plasticity in response to variations in prey availability [66, 67]. At a macro-spatial scale, the diet of Dupont’s Lark varied between Spain and Morocco, with greater frequency of grasshoppers (Acrididae, Orthoptera) in the diet of African birds. This may be related to habitat differences between countries, with Moroccan regions sampled dominated by alfa grass steppes, in contrast to the low scrub vegetation dominant in the Iberian steppes. Since the spatial distribution and abundance of many grasshopper species are closely linked to herbaceous vegetation [68, 69], which they use as food and shelter, it may be possible that the presence of grasshopper species is higher in the grass steppes inhabited by Dupont’s Lark in Morocco than in the Iberian shrub-steppes. Also, the phenology of grasshoppers may be an important factor in explaining the dietary differences found, since in Spain, the sampling time (early March-early June) was probably previous to the peak of adult grasshoppers’ activity, which in temperate regions usually starts in summer [70], while in Morocco, warmer conditions due to climate change have been reported to accelerate grasshopper development of species that show their peak in spring [71].

Dietary differences between continents might be also associated to morphological variations between Spanish and Moroccan populations. Feeding traits varied within the Dupont’s lark range probably in response to different natural selection forces acting in each part of the range [72]. Previous studies found that bill length and volume and tarsus length of Moroccan Dupont’s Larks were larger than in Spanish birds [24, 72], which likely increases accessibility to a broader range of prey species of different sizes [73]. These dietary differences were probably not sufficient to lead to a country-level diet differentiation, since at both sites individuals fed with similar frequency on other prey groups (beetles, lepidopterans, millipedes and spiders).

At country-level scale, the Dupont’s Lark population of Western Iberian Range (study site 5 in Fig 2B) appeared to be the most differentiated, primarily due to a notably higher frequency of lepidopteran consumption, especially from the families Noctuidae and Geometridae. The diet overlap analysis also supported this segregation of the Western Iberian Range, while the rest of the populations exhibited greater overlap in their diets.

Spatial variations in the diet composition of Dupont’s Lark may be influenced by landscape characteristics, which strongly shape the availability of insects and other arthropods, especially through features such as plant species richness or landscape heterogeneity [74]. Indeed, dietary differences were also observed between habitats, mainly due to the higher frequency of consumption of noctuid and geometrid lepidopterans by Dupont’s Lark individuals feeding in thyme-gorse shrublands. Most of the thyme-gorse shrublands are located in the study site Western Iberian Range (see Fig I-S2 and Table I-S2 of Appendix S2), where lepidopteran species of the genus *Agrotis* (Noctuidae), specially *Agrotis pierreti*, present their main Iberian populations [75, 76]. However, monitoring arthropod availability in occupied areas should be essential to better understand the dietary differences found, as well as to investigate prey selection throughout Dupont’s Lark distribution (see e.g., [36]). It is important to note that these results are derived from an unbalanced sample size among populations and habitats. Specifically, the Western Iberian Range population and the habitat named thyme-gorse shrublands had notably larger sample sizes than other populations and habitats, respectively. This discrepancy may have impacted our findings, as samples from those areas may have included taxa absent in samples from other sites due solely to the sample size effect. Consequently, interpretations should be treated with caution.

### Phenological variation

As hypothesized, we found differences in the diet composition of Dupont’s Lark between breeding and non-breeding seasons in Spain, primarily due to an increase in the frequency of consumption of grasshoppers of the family Acrididae during the non-breeding period. This variation may reflect changes in grasshopper availability and richness [8], as most acridid species in the Iberian Peninsula start their adult activity period at early summer, many of them reaching their maximum population sizes in September and October [77]. Although not significant, we also found temporal differences in the millipede consumption, as this arthropod group was not present in the non-breeding samples, period in which Mediterranean species reproduce and burrow until the adults emerge in spring [78]. Despite the differences, a high degree of dietary overlap was observed between the species’ phenological periods, since important arthropod groups in the diet of the species were predated during both periods with similar frequencies (e.g., Carabidae, Curculionidae, Tenebrionidae, Geometridae or Lycosidae), which might be related to the resident status of the species [79]. Future work should provide information on changes in prey availability in the study area between periods to better understand temporal variations in the diet of the species.

Temporal dietary changes have also been reported for other insectivorous bird species [8, 80], but to our knowledge, this study is the first to describe this pattern for a steppe passerine species. However, these results should be taken with caution due to the small sample size of the non-breeding period, so future work should incorporate a greater number of samples and extend the temporal coverage to the annual period. The only previous work on the non-breeding diet of Dupont’s Lark, based on the observation of four gizzard contents of specimens from the Ebro Valley, indicated the consumption of seeds of three plant genera in addition to insects [33]. The consumption of plant material in autumn and winter appears to be a common strategy in other steppe larks such as the Eurasian Skylark (*Alauda arvensis* [81, 82]. In this study, however, the plant component of the diet of Dupont’s Lark has not been analyzed, since the high diversity of plant MOTUs obtained with the marker 18S made it difficult to discriminate between environmental contamination, secondary consumption or items that were actively consumed by the birds [38, 83]. Nonetheless, detailed information on plant consumption might also be essential for a complete understanding of the dietary ecology of the species, and especially relevant for designing conservation strategies during the non-breeding season, when the plant fraction of the diet may acquire greater importance.

### Sexual variation

Sexual dietary differences have already been described for other bird species, including generalist medium-sized passerines such as the Black Wheatear (*Oenanthe leucura* [16] or the Hawfinch (*Coccothraustes coccothraustes* [67]). In contrast to our predictions, we revealed a high overlap between sexes in the diet of Dupont’s Lark during the breeding season, suggesting that foraging behavior and nutritional requirements are probably equivalents in both sexes. During the breeding season, females and males are expected to have different energetic demands due to differential reproductive roles [17], with Dupont’s Lark females implicated in nest-building, egg-laying and incubating, although both sexes are involved in food provisioning to nestlings [84]. This may result in females facing a stronger trade-off between self-maintenance and breeding [85]. A plausible explanation might be that abundance of resources in the immediate nest surroundings is higher than elsewhere, enabling females to prey on arthropod groups with similar frequencies to males’, although direct comparisons of arthropod availability around and far from the nest would be required. We also expected dietary differences between sexes due sex dimorphism in the Dupont’s Lark (males have a larger body size and a longer bill than females [23, 24]), which might help to reduce intraspecific competition for food resources [17]. Therefore, the absence of sexual diet differentiation suggests that food availability may not be a limiting factor during the breeding season in the Iberian steppes [86]. However, although both sexes preyed on the same arthropod groups, it may be possible that males and females fed on prey of different size [87], which cannot be evaluated with the method used in this study. Lack of dietary differences between males and females in birds with sexual size dimorphism have also been found in other studies [13, 88].

It is important to note that these conclusions are based on a small number of samples from females (*n* = 12), since the trapping method was inherently biased towards males [43]. Further research should aim to increase the sample size of females, as well as consider complementary methods able to determine the size of prey consumed by each sex, in order to draw more robust inferences about sexual variation in Dupont’s Lark diet. Nevertheless, our findings provide a first approximation into the diet of females Dupont’s larks and about the dietary patterns of the species, essential to integrate the female fraction of the population in conservation strategies.

### Metabarcoding considerations

In any metabarcoding study, the selection of the most appropriate primer set is critical and can affect the results obtained and the subsequent ecological interpretations [89]. Here, we used a multi-marker approach combining a universal marker (18S) with a specific arthropod marker (ZBJ), which allowed us to minimize the problems of taxonomic resolution and the potential taxonomic biases of the different markers [90]. The 18S amplified a broader taxonomic diversity than the ZBJ, but at the expense of higher taxonomic resolution, whereas the ZBJ probably biased detection in favor of Lepidoptera and Diptera over other arthropod taxa [38]. Although ZBJ usually provides identifications with high taxonomic resolution, we only identified 37% of the MOTUs at the species level, so dietary comparisons were performed at the order and family level. Therefore, dietary overlap and differences in prey composition for all the factors studied were conditioned by the level of identification reached. Had we been able to perform dietary comparisons at the species level, we would probably have detected greater differences and lower overlaps in our results.

Furthermore, metabarcoding is unable to discriminate life stages of consumed arthropods (adults, larvae or eggs), essential information for a comprehensive knowledge of the food ecology of Dupont’s Lark. Finally, our dietary study was based on presence/absence data rather than on read counts, since DNA-based methods remain subjected to a variety of factors (e.g. differential digestion rates of tissues, differences of copy numbers of marker genes between tissue types or prey taxa, PCR primer bias) that prevent to accurately correspond reads to the amount of each prey item consumed [39]. These and other considerations, including sample preservation and storage conditions, the incorporation of biological and technical replicates, or the use of negative and positive controls, should be taken into account when designing a DNA metabarcoding study to provide representative results of species’ diet.

## Conclusions and implications for Dupont’s Lark conservation

Metabarcoding represents a methodological advance for the study of insectivorous bird diets, providing greater accuracy for the identities and frequencies of prey taxa consumed compared to traditional techniques [38]. This study provides the first molecular insight into the diet of Dupont’s Lark across its European and Moroccan distribution range. The species fed on a wide range of arthropod prey, showing dietary differences between countries, populations and phenological periods, probably reflecting the plasticity and adaptability of the species to temporal and landscape differences in resource availability. Dietary plasticity can be important to this critically endangered bird [31], as it means that the species has certain ability to cope with unusual challenges such as food shortages [91]. However, arthropods are experiencing global declines in diversity and abundance in recent decades [92, 93], mostly due to habitat loss and degradation, and to agricultural intensification (pesticide usage, increased use of fertilizers and frequency of agronomic measures [94, 95]. The loss of arthropod biomass is expected to provoke cascading effects on food webs [92], raising concern about food limitations that could adversely affect the viability of insectivorous birds [96]. Furthermore, climate change is known to have the potential to alter synchrony between food availability and timing of reproduction in birds [97], which may have also important consequences for Dupont’s Lark populations. We also found that the diet of this species does not vary between sexes during the breeding season, suggesting similar food requirements and foraging habits of males and females, although additional research at a broader spatial scale and with a larger female sample size would be needed to further investigate sexual dietary patterns. Conservation and management measures focused on maintaining the quality of steppe habitat, including the promotion of traditional practices such as extensive sheep grazing [98], should be priority actions to ensure adequate availability and variety of essential food resources year-round for Dupont’s Lark and other insectivores of the steppe habitat.

## Acknowledgments

The authors wish to thank Israel Hervás, Miguel Muñoz, Ana Santos and Inmaculada Abril-Colón for their help during field and/or lab work. We also thank Vanessa A. Mata and Luís P. da Silva for their inestimable help with bioinformatic analysis. CP-G acknowledges support from Ministerio de Educación y Formación Profesional through the Beatriz Galindo Fellowship (Beatriz Galindo-Convocatoria 2020) and JZ acknowledges support from Ministerio de Universidades through the predoctoral FPU fellowship program.

## Supporting information

### Appendix S1. Bioinformatic pipeline description

### Appendix S2. Variation in diet composition between Spanish habitats. Fig I-S2. Location of Spanish sampling points classified by habitat

**Fig II-S2. Spider plot of the non-metric multidimensional scaling for arthropod prey families consumed by Dupont’s Lark across the Spanish habitats defined in this study.**

**Table I-S2. Number of Dupont’s Lark fecal samples per habitat in Spain and per population included in each habitat.**

**Table II-S2. Pairwise Pianka’s index values between habitats based on the frequency of arthropod families consumed by Dupont’s Lark.**

### Appendix S3. Prey items detected with the multi-marker approach and sample metadata

**Table I-S3. Prey items detected with the multi-marker approach in Dupont’s Lark fecal samples from Spain and Morocco.**

**Table II-S3. Sample metadata.**

### Appendix S4. Frequency of occurrence (%) of prey orders and families

**Table I-S4. Frequency of occurrence (%) of prey orders and families in the diet of Dupont’s Lark calculated for each country, sex, phenological period, Spanish population and Spanish habitat.**

### Appendix S5. MGLMs for differences in prey composition

**Table I-S5. Results of order-level and family-level multivariate generalized models (MGLMs) for the differences in diet composition between Spain and Morocco. Results of univariate tests are also displayed. Significant *P* values are indicated in bold.**

**Table II-S5. Results of order-level and family-level multivariate generalized models (MGLMs) for the differences in diet composition between Spanish populations. Results of the pairwise comparison and the univariate tests are also displayed. Significant *P* values are indicated in bold.**

**Table III-S5. Results of order-level and family-level multivariate generalized models (MGLMs) for the differences in diet composition between Spanish habitats. Results of the pairwise comparison and the univariate tests are also displayed. Significant *P* values are indicated in bold.**

**Table IV-S5. Results of order-level and family-level multivariate generalized models (MGLMs) for the differences in diet composition between the breeding and non-breeding seasons. Results of univariate tests are also displayed. Significant *P* values are indicated in bold.**

### Appendix S6. Phenological and sexual NMDS

**Fig I-S6. Spider plot of the non-metric multidimensional scaling for arthropod prey families consumed by Dupont’s Lark in the breeding and non-breeding seasons.**

**Fig II-S6. Spider plot of the non-metric multidimensional scaling for arthropod prey families consumed by Dupont’s Lark males and females in the breeding season.**

## References

1. Robbins CT. Wildlife feeding and nutrition, 4th edn. Academic Press, New York; 1993.

2. Durst SL, Theimer TC, Paxton EH, Sogge MK. Age, habitat, and yearly variation in the diet of a generalist insectivore, the Southwestern Willow Flycatcher. Condor. 2008;110(3): 514–525.

3. Begg CM, Begg KS, Du Toit JT, Mills MGL. Sexual and seasonal variation in the diet and foraging behaviour of a sexually dimorphic carnivore, the honey badger (*Mellivora capensis*). J Zool. 2003;260(3): 301–316.

4. Clare EL, Barber BR, Sweeney BW, Hebert PDN, Fenton MB. Eating local: influences of habitat on the diet of little brown bats (*Myotis lucifugus*). Mol Ecol 2011;20(8): 1772–1780.

5. Murray SW, Kurta A. Spatial and temporal variation in diet. In: Kurta A, Kennedy J, editors. The Indiana bat: Biology and management of an endangered species. Austin, TX: Bat Conservation International; 2002. pp. 182–192.

6. Pollard KA, Holland JM. Arthropods within the woody element of hedgerows and their distribution pattern. Agric For Entomol. 2006;8(3): 203–211.

7. Beck ML, Hopkins WA, Jackson BP. Spatial and temporal variation in the diet of Tree Swallows: implications for trace-element exposure after habitat remediation. Arch Environ Contam Toxicol. 2013;65(3): 575–587.

8. Davies SR, Vaughan IP, Thomas RJ, Drake LE, Marchbank A, Symondson WOC. Seasonal and ontological variation in diet and age-related differences in prey choice, by an insectivorous songbird. Ecol Evol. 2022;12: e9180.

9. Tournayre O, Leuchtmann M, Galan M, Trillat M, Piry S, Pinaud D, et al. eDNA metabarcoding reveals a core and secondary diets of the greater horseshoe bat with strong spatio-temporal plasticity. Environ DNA. 2021;3(1): 277–296.

10. Pyke GH, Pulliam HR, Charnov EL. Optimal foraging: a selective review of theory and tests. Q Rev Biol. 1977;52: 137–154.

11. Ramos R, Ramírez F, Carrasco JL, Jover L. Insights into the spatiotemporal component of feeding ecology: an isotopic approach for conservation management sciences. Divers Distrib. 2011;17(2): 338–349.

12. Wellicome TI, Danielle Todd L, Poulin RG, Holroyd GL, Fisher RJ. Comparing food limitation among three stages of nesting: supplementation experiments with the burrowing owl. Ecol Evol. 2013;3(8): 2684–2695.

13. Navarro J, Kaliontzopoulou A, González-Solís J. Sexual dimorphism in bill morphology and feeding ecology in Cory’s shearwater (*Calonectris diomedea*). Zool. 2009;112(2): 128–138.

14. Zurdo J, Barrero A, da Silva LP, Bustillo-de la rosa D, Gómez-Catasús J, Morales MB, et al. Dietary niche overlap and resource partitioning among six steppe passerines of Central Spain using DNA metabarcoding. Ibis. 2023;165(3): 905–923.

15. Bravo C, Ponce C, Bautista LM, Alonso JC. Dietary divergence in the most sexually size-dimorphic bird. Auk. 2016;133(2): 178–197.

16. da Silva LP, Mata VA, Lopes PB, Lopes RJ, Beja P. High-resolution multi-marker DNA metabarcoding reveals sexual dietary differentiation in a bird with minor dimorphism. Ecol Evol. 2020;10(19): 10364–10373.

17. Catry P, Phillips RA, Croxall JP. Sexual segregation in birds: patterns, processes and implications for conservation. In: Ruckstuhl KE, Neuhaus P, editors. Sexual segregation in vertebrates: ecology of the two sexes. Cambridge, UK: Cambridge University Press; 2005. pp. 351–378.

18. Bolnick DI, Doebeli M. Sexual dimorphism and adaptive speciation: two sides of the same ecological coin. Evol. 2003;57: 2433–2449.

19. Ruckstuhland K, Neuhaus P. Sexual Segregation in Vertebrates: Ecology of the Two Sexes. New York: Cambridge University Press; 2006.

20. García JT, Suárez F, Garza V, Justribó JH, Oñate JJ, Hervás I, et al. Assessing the distribution, habitat, and population size of the threatened Dupont’s Lark *Chersophilus duponti* in Morocco: Lessons for conservation. Oryx. 2008;42: 592.

21. BirdLife International. Species factsheet: *Chersophilus duponti*; 2023. Available from http://datazone.birdlife.org/species/factsheet/duponts-lark-chersophilus-duponti on 07/10/2023.

22. Traba J, Pérez-Granados C, Serrano D. Alondra ricotí. Cherosphilus duponti. In: López-Jiménez J, editor. Libro Rojo de las Aves de España. SEO/BirdLife, Madrid; 2021. pp. 321–329.

23. Vögeli M, Serrano D, Tella JL, Méndez M, Godoy JA. Sex determination of Dupont’s lark *Chersophilus duponti* using molecular sexing and discriminant functions. Ardeola. 2007;54(1): 69–79.

24. García-Antón A, Garza V, Traba J. Climate, isolation and intraspecific competition affect morphological traits in an endangered steppe bird, the Dupont’s Lark *Chersophilus duponti*. Bird Study. 2018;65(3): 373–384.

25. Garza V, Suárez F, Herranz J, Traba J, García de la Morena EL, Morales MB, et al. Home range, territoriality and habitat selection by the Dupont’s lark *Chersophilus duponti* during the breeding and postbreeding periods. Ardeola. 2005;52: 133–146.

26. Laiolo P, Tella JL. Fate of unproductive and unattractive habitats: Recent changes in Iberian steppes and their effects on endangered avifauna. Environ Conserv. 2006;33(3): 223–232.

27. Traba J, Morales MB. The decline of farmland birds in Spain is strongly associated to the loss of fallowland. Sci Rep. 2019;9: 9473.

28. Viñuela J, García JT, Suárez F. Marked range regression and possible alteration of distribution of the Dupont’s Lark *Chersophilus duponti* in Tunisia: Conservation consequences of vanishing alfa grass *Stipa tenacissima* steppes in North Africa. Diversity. 2023;15: 549.

29. Oñate JJ, Suárez F, Calero-Riestra M, Justribó JH, Hervás I, de la Morena ELG, et al. Responses of bird communities to habitat structure along an aridity gradient in the steppes north of the Sahara. Diversity. 2023;15: 737.

30. García-Antón A, Garza V, Traba J. Connectivity in Spanish metapopulation of Dupont’s lark may be maintained by dispersal over medium-distance range and stepping stones. PeerJ. 2021;9: e11925.

31. Reverter M, Pérez-Granados C, López-Iborra GM, García-Mellado A, Aledo-Olivares E, Alcántara M, et al. Range contraction and population decline of the European Dupont’s Lark population. Diversity. 2023;15: 928.

32. Cramp S. Handbook of the Birds of Europe, Middle East and North Africa: The Birds of the Western Palearctic. Volume V. Tyrant Flycatchers to Thrushes: 1063. Oxford: Oxford University Press; 1988.

33. Aragüés A. La alondra de Dupont (*Chersophilus duponti*) en Monegros. Boletín de la S.E.A. 1999;24: 196–198.

34. Herranz J, Yanes M, Suárez F. Primeros datos sobre la dieta de pollos de Alondra de Dupont, *Chersophilus duponti*, en la Península Ibérica. Ardeola. 1993;40: 77–79.

35. Zurdo J, Gómez-López P, Barrero A, Bustillo-de la Rosa D, Gómez-Catasús J, Reverter M, et al. Selecting the best: Interspecific and age-related diet differences among sympatric steppe passerines. Avian Res. 2023;14: 100151.

36. Zurdo J, Reverter M, Barrero A, Bustillo-de la Rosa D, Gómez-Catasús J, Pérez-Granados C, et al. Prey choice in insectivorous steppe passerines: New insights from DNA metabarcoding. Glob Ecol Conserv. 2023;48: e02738.

37. Pompanon F, Deagle BE, Symondson WOC, Brown DS, Jarman SN, Taberlet P. Who is eating what: Diet assessment using next generation sequencing. Mol Ecol 2012;21: 1931–1950.

38. da Silva LP, Mata VA, Lopes PB, Pereira P, Jarman SN, Lopes RJ, et al. Advancing the integration of multimarker metabarcoding data in dietary analysis of trophic generalists. Mol Ecol Resour. 2019;19: 1420–1432.

39. Hoenig BD, Snider AM, Forsman AM, Hobson KA, Latta SC, Miller ET, et al. Current methods and future directions in avian diet analysis. Ornithology. 2022;139(1): ukab077.

40. Suárez F. La alondra ricotí (Chersophilus duponti). Dirección General para la Biodiversidad. Ministerio de Medio Ambiente y Medio Rural y Marino Medio Rural y Marino, Madrid; 2010.

41. Bustillo-de La Rosa D, Traba J, Calero-Riestra M, Morales MB, Barrero A, Viñuela J, et al. Recent changes in genetic diversity, structure, and gene flow in a passerine experiencing a rapid population decline, the Dupont’s Lark (*Chersophilus duponti*). Diversity. 2022;14(12): 1120.

42. García-Antón A, Garza V, Hernández Justribó J, Traba J. Factors affecting Dupont’s Lark distribution and range regression in Spain. PLOS ONE. 2019;14: e0211549.

43. Barrero A, Gómez-Catasús J, Pérez-Granados C, Bustillo-de la Rosa D, Traba J. Conspecific density and habitat quality drive the defence and vocal behaviour of a territorial passerine. Ibis. 2023.

44. Cabodevilla X, Gómez-Moliner BJ, Abad N, Madeira MJ. Simultaneous analysis of the intestinal parasites and diet through eDNA metabarcoding. Integr Zool. 2023;18: 399–413.

45. Zeale MRK, Butlin RK, Barker GLA, Lees DC, Jones G. Taxon-specific PCR for DNA barcoding arthropod prey in bat faeces. Mol Ecol Resour. 2011;11: 236–244.

46. Boyer F, Mercier C, Bonin A, Le Bras Y, Taberlet P, Coissac E. OBITools: A unix-inspired software package for DNA metabarcoding. Mol Ecol Resour. 2016;16: 176– 182.

47. Rognes T, Flouri T, Nichols B, Quince C, Mahé F. VSEARCH: A versatile open source tool for metagenomics. PeerJ. 2016;4: 1–22.

48. Mahé F, Rognes T, Quince C, de Vargas C, Dunthorn M. Swarm v2: Highly-scalable and high-resolution amplicon clustering. PeerJ. 2015;3: 1–12.

49. Frøslev TG, Kjøller R, Bruun HH, Ejrnæs R, Brunbjerg A.K, Pietroni C, et al. Algorithm for post-clustering curation of DNA amplicon data yields reliable biodiversity estimates. Nat Commun. 2017;8: 1188.

50. Drake LE, Cuff JP, Young RE, Marchbank A, Chadwick EA, Symondson WOC. An assessment of minimum sequence copy thresholds for identifying and reducing the prevalence of artefacts in dietary metabarcoding data. Methods Ecol Evol. 2021;13: 1–17.

51. Lamb PD, Hunter E, Pinnegar JK, Creer S, Davies RG, Taylor MI. How quantitative is metabarcoding: A meta-analytical approach. Mol. Ecol. 2019;28(2): 420–430.

52. R Core Team. R: A Language and Environment for Statistical Computing. R Foundation for Statistical Computing, Vienna, Austria; 2023. https://www.R-project.org/

53. Wang Y, Naumann U, Eddelbuettel D, Wilshire J, Warton D. mvabund: Statistical Methods for Analysing Multivariate Abundance Data. R package version 4.2.1; 2022. https://CRAN.R-project.org/package=mvabund

54. Oksanen J, Simpson G, Blanchet F, Kindt R, Legendre P, Minchin P, et al. vegan: Community Ecology Package. R package version 2.6-4; 2022. https://CRAN.R-project.org/package=vegan

55. Pianka ER. The structure of lizard communities. Annu Rev Ecol Evol Syst. 1973;4: 53–74.

56. Gotelli NJ, Hart EM, Ellison AM. EcoSimR: Null model analysis for ecological data. R package version 0.1.0; 2015. http://github.com/gotellilab/EcoSimR

57. Lawlor LR. Structure and stability in natural and randomly constructed model ecosystems. Am Nat. 1980;116: 394–408.

58. Aguirre JL, Talabante C, Aparicio A, Peinado M. Phytosociological, structural and conservation analysis of the habitats of Dupont’s lark in Europe: a phytosociological survey applied to the conservation of an endangered species. Plant Biosyst. 2017;152(5): 953–970.

59. Bedrosian G, Watson JW, Steenhof K, Kochert MN, Preston CR, Woodbridge B, et al. Spatial and temporal patterns in Golden Eagle diets in the Western United States, with implications for conservation planning. J Raptor Res. 2017;51(3): 347–367.

60. Bureš S, Weidinger K. Sources and timing of calcium intake during reproduction in flycatchers. Oecologia. 2003;137: 634–647.

61. Razeng E, Watson DM. Nutritional composition of the preferred prey of insectivorous birds: popularity reflects quality. J Avian Biol. 2015;46: 89–96.

62. Talabante C, Aparicio A, Aguirre JL, Peinado M. Avances en el estudio de la alimentación de adultos de Alondra ricotí (*Chersophilus duponti*) y la importancia de los escarabajos coprófagos. I Workshop nacional de la Alondra Ricotí: Estrategias futuras. Estación Ornitológica de Padul (EOP), Granada; 2015.

63. Diarra SS. Utilisation of snail meal as a protein supplement in poultry diets. Poult Sci J. 2015;71(3): 547–554.

64. Guennoun FZ, Mostakim L, Ghamizi M. Biodiversity assessment of terrestrial snails (Mollusca, Gastropoda) of Essaouira’ dunes of Morocco: testing factors affecting the distribution of terrestrial molluscs. Appl Ecol Environ Res. 2023;21(3).

65. Yanes M, Suárez F, Manrique J. La cogujada montesina, *Galerida theklae*, como depredador del caracol *Otala lactea*: Comportamiento alimenticio y selección de presa. Ardeola. 1991;38(2): 297–303.

66. Shin D-M, Yoo J-C, Jeong D-M. Spatial variation of Eurasian Eagle-Owl diets in wetland and non-wetland habitats in West-Central Korea. J Raptor Res. 2013;47(4): 400–409.

67. Stenhouse EH, Bellamy P, Kirby W, Vaughan IP, Drake LE, Marchbank A, et al. Multi-marker DNA metabarcoding reveals spatial and sexual variation in the diet of a scarce woodland bird. Ecol Evol. 2023;13: e10089.

68. Badenhausser I, Gross N, Cordeau S, Bruneteau L, Vandier M. Enhancing grasshopper (Orthoptera: Acrididae) communities in sown margin strips: the role of plant diversity and identity. Arthropod-Plant Interact. 2015;9(4): 333–346.

69. Wei S, Liu X, McNeill MR, Wang Y, Sun W, Tu X, et al. Identification of spatial distribution and drivers for grasshopper populations based on geographic detectors. Ecol Indic. 2023;154: 110500.

70. Aguirre-Segura A, Barranco P. Orden Orthoptera. Ibero Diversidad Entomológica. 2015;46: 1–13.

71. Imene BS, Abboud H, Daniel P. Phenology of early-season and mid-season grasshoppers shows contrasted responses toward climatic variations in an arid area. Intern J Zool. 2019;5(1): 22–32.

72. García JT, Suárez F, Garza V, Calero-Riestra M, Hernández J, & Pérez-Tris J. Genetic and phenotypic variation among geographically isolated populations of the globally threatened Dupont’s lark *Chersophilus duponti*. Mol Phylogenetics Evol. 2008;46(1): 237–251.

73. Leisler B, Winkler H. Evolution of island warblers: beyond bills and masses. J Avian Biol. 2015:46(3): 236–244.

74. Bonari G, Fajmon K, Malenovský I, Zelený D, Holuša J, Jongepierová I, et al. Management of semi-natural grasslands benefiting both plant and insect diversity: The importance of heterogeneity and tradition. Agric Ecosyst Environ. 2017;246: 243–252.

75. Calle JA. Noctuidos españoles. Ministerio de Agricultura, Pesca y Alimentación. Madrid; 1982.

76. Magro R, Jambrina J. Catálogo razonado de los Lepidoptera de Castilla y León, España (Parte III) (Lepidoptera: Notodontidae, Euteliidae, Noctuidae). SHILAP Rev Lepidopt. 2014;42: 173–212.

77. Moyano L. Estudio y seguimiento de la fauna de Orthoptera de un entorno natural sometido a un programa de restauración ecológica en el sur de la Península Ibérica. Doctoral Thesis. Universidad de Córdoba. 2014.

78. David JF, Coulis M. Millipedes faced with drought: the life cycle of a Mediterranean population of *Ommatoiulus sabulosus* (Linnaeus) (Diplopoda, Julida, Julidae). Zookeys. 2015;510: 115–24.

79. Gómez-Catasús J, Barrero A, Garza V, Traba J. Alondra ricotí – Chersophilus duponti. In: Salvador A, Morales MB, editors. Enciclopedia Virtual de los Vertebrados Españoles. Museo Nacional de Ciencias Naturales, Madrid; 2016. http://www.vertebradosibericos.org/

80. McClenaghan B, Nol E, Kerr KCR. DNA metabarcoding reveals the broad and flexible diet of a declining aerial insectivore. Auk 2019;136: 1–11.

81. Suárez F, Hervás I, Herranz J. Las alondras de España peninsular. Madrid: Dirección General para la Biodiversidad, Ministerio de Medio Ambiente y Medio Rural y Marino; 2009.

82. Eraud C, Cadet E, Powolny T, Gaba S, Bretagnolle F, Bretagnolle V. Weed seeds, not grain, contribute to the diet of wintering skylarks in arable farmlands of Western France. Eur J Wildl Res. 2014;61(1): 151–161.

83. Tercel MP, Symondson WOC, Cuff JP. The problem of omnivory: A synthesis on omnivory and DNA metabarcoding. Mol Ecol. 2021;30: 2199–2206.

84. Pérez-Granados C, López-Iborra GM, Garza V, Traba J. Breeding biology of the endangered Dupont’s Lark *Chersophilus duponti* in two separate Spanish shrub-steppes. Bird Study. 2017;64(3): 328–338.

85. Mainwaring MC, Hartley IR. Causes and consequences of differential growth in birds. Adv Stud Behav. 2012;44: 225–277.

86. Barrero A, Ovaskainen O, Traba J, Gómez-Catasús J. Co-occurrence patterns in a steppe bird community: insights into the role of dominance and competition. Oikos. 2023; e09780.

87. Blondel J, Perret P, Anstett MC, Thebaud C. Evolution of sexual size dimorphism in birds: test of hypotheses using blue tits in contrasted Mediterranean habitats. J Evol Biol. 2002;15(3): 440–450.

88. Perrig PL, Lambertucci SA, Alarcón PAE, Middleton AD, Padró J, Plaza PI, et al. Limited sexual segregation in a dimorphic avian scavenger, the Andean condor. Oecologia. 2021;196(1): 77–88.

89. Alberdi A, Aizpurua O, Bohmann K, Gopalakrishnan S, Lynggaard C, Nielsen M, et al. Promises and pitfalls of using high-throughput sequencing for diet analysis. Mol Ecol Resour. 2019;19(2): 327–348.

90. Forsman AM, Hoenig BD, Gaspar SA, Fischer JD, Siegrist J, Fraser K. Evaluating the impacts of metabarcoding primer selection on DNA characterization of diet in an aerial insectivore, the Purple Martin. Ornithology. 2022;139(1): ukab075.

91. Ducatez S, Sol D, Sayol F, Lefebvre L. Behavioural plasticity is associated with reduced extinction risk in birds. Nat Ecol Evol. 2020;4(6): 788–793.

92. Hallmann CA, Sorg M, Jongejans E, Siepel H, Hofland N, Schwan H, et al. More than 75 percent decline over 27 years in total flying insect biomass in protected areas. PLOS ONE. 2017;12(10): e0185809.

93. Wagner DL, Grames EM, Forister ML, Berenbaum MR, Stopak D. Insect decline in the Anthropocene: death by a thousand cuts. PNAS 2021;118(2): e2023989118.

94. Fox R. The decline of moths in Great Britain: a review of possible causes. Insect Conserv Divers. 2013;6(1): 5–19.

95. Reverter M, Gómez-Catasús J, Barrero A, Traba J. Crops modify habitat quality beyond their limits. Agric Ecosyst Environ. 2021;319: 107542.

96. Grames EM, Montgomery GA, Youngflesh C, Tingley MW, Elphick CS. The effect of insect food availability on songbird reproductive success and chick body condition: Evidence from a systematic review and meta-analysis. Ecology Letters, 26(4), 658–673. Appl Ecol Environ Res. 2023;21(3): 1957–1978.

97. Visser ME, Both C. Shifts in phenology due to global climate change: the need for a yardstick. Proc R Soc Lond B Biol Sci. 2005;272: 2561–2569.

98. Gómez-Catasús J, Reverter M, Bustillo-de la Rosa D, Barrero A, Pérez-Granados C, Zurdo, J, et al. Moderate sheep grazing increases arthropod biomass and habitat use by steppe birds. Agric Ecosyst Environ. 2023;354: 108556.

